# Individual variability in the human connectome maintains selective cross-modal consistency and shares microstructural signatures

**DOI:** 10.1101/2021.04.01.438129

**Authors:** Esin Karahan, Luke Tait, Ruoguang Si, Ayşegül Özkan, Maciek Szul, Jiaxiang Zhang

## Abstract

Individuals are different in their behavioural responses and cognitive abilities. Neural underpinnings of individual differences are largely unknown. Here, by using multimodal imaging data including diffusion MRI, functional MRI and MEG, we show the consistency of interindividual variation of connectivity across modalities. We demonstrated that regional differences in individual variability of structural and functional connectomes is characterized by higher variability in association cortices and lower variability in sensory and visual cortices. This pattern is consistent across all modalities at varying degrees as shown by significant alignment between functional and structural connectome variabilities at several clusters of brain regions. Variability in connectivity is associated with cortical myelin content and microstructural properties of connections. Our findings contribute to understanding of individual differences in functional and structural organization of brain and facilitate fingerprinting applications.

## 1. Introduction

Individuals are unique as determined by their unique copy of DNA. Interplay between environment and genetic factors gives rise to differences in behavior and cognition. Finding neural underpinnings of individual differences is one of the greatest endeavors in neuroscience research. Similar to DNA fingerprinting, by encoding connections in the brain neural fingerprinting could reliably identify individuals at anatomical [1], structural [2, 3], functional [4] and electrophysiological levels [5]. Identification of individual differences in healthy population and linking them to cognitive variability is helpful to associate differences with disease population.

MR based imaging methods have been used in quantifying individual differences due to high spatial resolution [6, 7]. Structural connectome is derived from strength of connections between brain regions estimated from tractography with diffusion MRI (dMRI). Functional connectome is a statistical measure of co-activation of brain region pairs based on fMRI. Interindividual variability in functional connectome has been shown to be associated with cortical surface expansion, anatomical variability, and long-range integration. Weak correlation was found between structural and functional connectome variability [7]. Research on individual differences brings three unresolved issues. First, fMRI derived functional connectome cannot capture fast dynamics and oscillations of brain activity in milliseconds resolution [8]. On the other hand, MEG can measure postsynaptic activity of pyramidal neurons located perpendicular to cortical surface at higher temporal resolution with a limited spatial scale [9]. Since both modalities capture complementary aspects of functional connectivity, inter-individual differences based in these methods can discover variability in both stable and transient networks. Second, alignment between modalities in terms of individual variability is largely unknown. Different imaging modalities could give varying information due to underlying genuine neurobiological variability or due to limits of modality itself. Third, relation of variability in macroscale metrics to microscopic tissue architecture is not known. Microstructural properties of white matter tracts predict cross-subject variance in functional connectivity in interhemispheric connections of homotopic regions [10].

We addressed these questions by using a multimodal imaging data including dMRI, fMRI and MEG. MEG based functional connectome is calculated on envelope of time series of cortical sources. We generated structural and functional connectomes on a recent multimodal atlas [11] for comparison. Since areal size effects structural connectome at great extent [12] we subparcellated atlas to accommodate for large differences between small and big parcels. We tested multimodal alignment by calculating individual variability between pairs of individuals of structural and functional connectomes on clusters based on multimodal atlas.

Microstructure is acquired at both gray matter and white matter level. To relate with microstructure of gray matter, we measured quantitative R1 values as a proxy to myelination on an independent cohort. White matter microstructure is calculated from dMRI based on both conventional DTI and biophysical models. We compared whether gray matter microstructure of a region and white matter microstructure profile of white matter can predict variabilities in functional connectomes.

Our results demonstrated that all imaging modalities reflect non-homogenous individual variability pattern across brain. Higher variability in association cortices and lower variability in unimodal cortices is a common property for all modalities at varying degrees. Gray matter and white matter microstructure partially explain multimodal individual variability.

## 2. Methods

### 2.1. Participants

64 healthy participants were recruited from the Cardiff University School of Psychology participant panel (49 females, age range 18–35 years; mean age 21.1 ±2.94 years). Cohort 1 included 29 participants (20 females, age range 18-28 years; mean age 20.76 ±2.14 years). All participants in Cohort 1 underwent a 3T MRI session, and 28 participants in Cohort 1 completed two further MEG sessions. Cohort 2 included 35 participants (29 females, age range 18-35 years; mean age 21.34 ±3.5 years), and all completed a 7T structural MRI session. No participant reported a history of neurological or psychiatric illness. The study was approved by the Cardiff University School of Psychology Research Ethics Committee. All participants gave written informed consent.

### 2.2. MRI data acquisition for Cohort 1

Whole-brain, multi-shell, diffusion-weighted images (DWI) were acquired from all participants in Cohort 1, using a Siemens 3T Connectom MRI scanner (Siemens Medical Systems) at the Cardiff University Brain Research Imaging Centre (CUBRIC). The spin-echo echoplanar imaging (EPI) pulse sequence used a HARDI protocol (echo time 59 ms, repetition time 3000 ms, voxel size 2×2×2 mm). Diffusion sensitizing gradients were applied in 20 isotropic directions at *b*-values of 200 and 500 s/mm^2^, in 30 isotropic directions at a b-value of 1200 s/mm^2^ and in 61 isotropic directions at b-values of 2400, 4000, 6000 s/mm^2^. Thirteen volumes with no diffusion weighting (*b* = 0 s/mm^2^) interleaved across the sequence were also acquired. To correct for susceptibility induced distortions, three images at *b* = 0 s/mm^2^ and 30 diffusion directions at *b* = 1200 s/mm^2^ were acquired with the opposite phase encoding direction. Participants also underwent high-resolution T1-weighted magnetization prepared rapid gradient echo scanning (MP-RAGE: echo time 3.06 ms; repetition time 2250 ms sequence, flip angle 9°, field of view=256×256 mm, acquisition matrix 256×256, voxel size 1×1×1 mm).

Whole brain, BOLD-sensitive T2*-weighted EPI images were acquired using a multi-band protocol (echo time 35 ms, repetition time 1500 ms, voxel size 2 × 2 × 2 mm, multi-band factor 3, flip angle 70°, AC-PC alignment with a posterior-down tilt). 315 volumes were acquired with an interleaved order. During the scan, participants were instructed to rest with their eyes open and fixate on a red dot with a grey background, presented through a back projector.

### 2.3. MEG data acquisition for Cohort 1

Whole head MEG recordings were acquired in a magnetically shielded chamber, using a 275-channel CTF radial gradiometer system (CTF Systems, Canada) at a sampling rate of 1200 Hz. One sensor was turned off during recording due to excessive noise. Additional 29 reference channels were recorded for noise cancellations and the primary sensors were analyzed as synthetic third-order gradiometers [13]. Continuous horizontal and vertical bipolar electro-oculograms (EOG) were recorded to monitor blinks and eye movements. Participants seated comfortably in the MEG chair and their head was supported with a chin rest to minimise head movement. For MEG/MRI co-registration, head shape with the position of coils was digitized using a Polhemus FASTRAK (Colchester, Vermont). Participants were instructed to rest with their eyes open and fixate on a red dot with a grey background, presented through a back projector. Each recording session lasted approximately 8 minutes. 28 participants in Cohort 1 underwent two resting-state MEG sessions on different days (1 to 8 days between two sessions). For all MEG analyses, we therefore analyzed data from those 28 participants.

### 2.4. Cortical reconstruction

Freesurfer (version 5.3.0, http://surfer.nmr.mgh.harvard.edu) was used to process T1-weighted MP-RAGE images, including motion correction, intensity normalization, skull-stripping, white-matter segmentation, tessellation, smoothing, inflating and spherical registration [14]. After pre-processing, the surface of grey matter/white matter boundary was generated, together with inner skull, scalp and pial images. Conformed and intensity normalized T1-weighted image was registered to the mean non-diffusion image (*b* = 0 s/mm^2^) by using a boundary-based rigid body registration with six degrees of freedom [15]. For each participant, the forward and inverse transformation matrices between the native DWI space and T1 space were used for subsequent co-registration and tractography analyses.

#### 2.4.1. Parcellation

Multimodal Human Connectome Project based (HCP-MMP) atlas was used for the parcellation of the brain surface into 360 regions [11]. Since hippocampal parcellations were not successful in all subjects, we discarded regions labeled as hippocampus in both hemispheres leading to 358 regions in total. Size of the areas of those regions were highly variable (122 mm^2^ smallest region to 3198 mm^2^ largest region) in original HCP-MMP atlas. To discard confounding effects of nonhomogeneous areal size across brain on the subsequent interindividual variance analysis, we subdivided larger regions in HCP-MMP atlas by using *”mris_divide_parcellation”* command from Freesurfer. The number of subdivisions per region was determined to ensure that variability of areal size was minimal (average largest region size was 454 mm^2^). We manually checked and corrected for small and large subdivisions in each region. Final number of region of interests (ROIs) was 664. We generated tractography according to original 358 regions, but calculated structural connectome based on 664 ROIs.

Subject-level parcellation of subdivided HCP-MMP atlas was created by registering the annotation files in the fsaverage space to the subject space and later sampling surface into volume. Cortical parcellation volume per subject was co-registered to native DWI space without resampling and was dilated by assigning closest white matter voxels to the gray matter regions. Assigning was based on searching grey matter voxels in the cube of size 5×5×5mm with white matter voxel at the center, sorting the number of grey matter voxels that belong to a region and calculating the distance between white matter voxel and the grey matter region. Finally, if the distance was less than 1.73 mm, white matter voxel was assigned to that region.

White matter mask was created based on native space registered Freesurfer segmentation images that includes white matter and dilated gray matter/white matter boundary which was used in tractography.

### 2.5. DWI data preprocessing

DWI data were converted from DICOM to NIfTI format using dcm2nii. The images were skull-stripped using FSL BET and denoised for thermal noise with MP-PCA method [16] using denoise tool from MRTrix (MRTrix 3), [17]. Following drift correction [18], images were corrected for susceptibility induced distortions, eddy currents and head motion using FSL eddy and topup functions (FSL 6.0.1). After correcting for gradient nonlinearity [19] and Gibbs ringing artefacts [20], mean non-diffusion image (*b* = 0 s/mm^2^) was formed.

### 2.6. Tractography

Input response functions for CSF, GM and single fiber WM were estimated from the multishell DWI data [21]. Using these three functions, fibre orientation distributions were estimated with multi-tissue constrained spherical deconvolution [22] for each tissue compartment in all voxels. Finally, multi-tissue fiber orientation functions were normalized with *mtnormalise* tool from MRTrix to enable multisubject comparison.

To generate seed masks for tractography, first we resampled left and right hemisphere surfaces of each subject into volume to create white matter/gray matter boundary image and registered to native DWI space by using transformation matrix created in previous step. Then, binarized WM/GM boundary image was multiplied with subject based dilated HCP-MMP atlas image to create a labeled WM/GM boundary image. Subcortical segmentation created in the recon-all procedure was added to this labeled image to consider these regions in tractography. In total 375 seed masks (358 cortical, 17 subcortical) were created.

A probabilistic algorithm based on the second-order integration over fiber orientation was used to calculate region to region tractography [23] (Tournier et al., 2010). Seed masks were limited to the HCP-MMP atlas parcellations on the dilated GM/WM boundary to avoid partial volume effects. Target masks were defined as the rest of the GM/WM boundary and subcortical regions except seed mask. Tractography was constrained to the white matter. A fixed number of streamlines per seed-voxel (200) was generated [24]. Minimum length of streamlines was determined as 10mm. To optimize the computational power of the tractography, larger seed masks were divided into smaller masks where target masks were kept the same [24] and tractography was performed in parallel.

#### 2.6.1. Streamline Trimming

Streamlines that did not end in the target mask or transverse WM were discarded. Furthermore, to eliminate outlier streamlines, outlier removal based on clustering was implemented similar to [25]. For region to region tracks, streamlines were sampled along their length, grouped into clusters according to Euclidean distance to each other and new clusters were formed if a streamline was more distant than predetermined threshold. Clusters with a smaller number of elements were discarded. After testing for several values, we used 5 for the distance threshold.

### 2.7. Structural Connectome

We generated structural connectome from 664 ROIs of subparcellated HCP-MMP atlas. To upsample from original HCP-MMP atlas-based seed masks to our newly generated subparcellated HCP-MMP atlas, we regrouped streamlines according to their seed and target voxels based on subparcellated atlas. Finally, structural connectome was extracted by counting the number of valid streamlines between each pair of ROIs. ROI to ROI connections that had less than 50 streamlines are discarded. We calculated connection probability of each ROI by dividing the number of streamlines between two ROIs with the total number of streamlines that were emitted by this particular ROI. Note that this operation resulted in non-symmetric structural connectome.

### 2.8. Microstructural Models

From the preprocessed DWI data, we calculated fractional anisotropy (FA), mean diffusivity (MD), radial diffusion (RD) and angular diffusion (AD) by fitting the b=0, 500 and 1200 s/mm^2^ shell data to DTI m odel. We used data from lower b values since DTI model is based on hindered diffusion in the extra-axonal space which is more sensitive to lower b values [26].

Same dataset was fitted to NODDI model using the NODDI MATLAB Toolbox v1.0.1 to calculate volume fraction of the intracellular compartment (ICVF) and orientation dispersion index (ODI). NODDI models brain tissue in three compartments comprising intracellular space, extracellular space, and CSF [27]. By this way, restricted diffusion perpendicular to neurites and unhindered diffusion along them is explicitly modeled. ICVF quantifies the volume of compartment that contains the axons and dendrites, whereas ODI represents angular variation in neurite orientation.

We calculated restricted diffusion signal fraction maps (FR) by fitting CHARMED [28] which characterizes white matter in restricted and hindered compartments. CHARMED is more sensitive to local fiber orientation with respect to standard DTI model thus giving a better estimate of restricted diffusion of intra-axonal water molecules.

### 2.9. Connectome Microstructure

We calculated 7 microstructural measures including four from DTI model: FA, MD, RD and AD, two from NODDI model: IVCF and ODI and finally FR from CHARMED model on nondiffusion DWI mask. All microstructural measures are sampled along the streamlines by using *tcksample* function from MRTrix [17]. We took the median value of measures along and across streamlines to characterize microstructural properties of region to region connections. By this way, we had 664 × 664 × 7 microstructural connectomes for each individual where connections that were discarded in structural connectome were not included. We converted microstructural measures to zscore per subject to avoid scale differences between measures and subjects.

Since microstructural measures are not completely separable and contain complementary information [29, 30], we combined microstructural measures to maximize the explained variance.

For this analysis, we vectorized lower triangle of microstructural matrices and concatenated along subject dimension to construct data matrix *subjects*(*n* = 29) · *connections*(*n* = 220116) × *microstrcuture*(*n* = 7). We applied principal component analysis (PCA) to find the orthogonal components that explain the 90%of variance along the microstructural dimension.

### 2.10. fMRI preprocessing

All resting state fMRI (rsfMRI) images were processed through the Connectome Computation System (CCS) [31] which integrates Freesurfer [32], FSL [33] and AFNI [34] to project 3D rsfMRI images onto 2D cortical surface. Projection on cortical surface can increase the test-retest reliability of rsfMRI analyses [35]. The main steps of functional preprocessing in CCS included (1) signal equilibration by removing the first 10 seconds of each scan, (2) slice timing correction, (3) 3D motion correction, (4) 4D global mean-based intensity normalization, (5) regressing out the white matter, cerebrospinal fluid and motion parameters, (6) band-pass temporal filtering (0.01–0.1 Hz), (7) removal of linear and quadratic trends, (8) co-registration between individual functional and anatomical images using a rigid boundary-based transformation (BBR) algorithm [15], and (10) projection of functional images onto the fsaverage5 cortical surfaces in the standard MNI space (10,242 vertices per hemisphere and 4 mm inter-vertex gap on average) [36].

### 2.11. fMRI based functional connectome

The rsfMRI functional connectivity matrices were calculated on the subparcellated HCP-MMP atlas. A summary ROI-level time series was obtained by averaging signals within the region. Finally, functional connectivity was computed by Pearson correlating time series data between every pair of ROIs, resulting in 664 × 664 FC matrices for each subject. We ignored connections that did not have significant correlation and later thresholded FC matrices to obtain approximately 10% edge density for each subject by removing weakest connections [36].

### 2.12. MEG preprocessing

MEG data preprocessing followed previously described pipelines [37]. Continuous raw MEG data was imported to Fieldtrip [38], downsampled to 256 Hz, bandpass filtered at 1-100 Hz (4th order two-pass Butterworth filter). Data was subsequently notch filtered at 50 and 100 Hz to remove line noise. Visual and cardiac artifacts were removed using ICA decomposition, using the *fastica* algorithm [39]. Identification of visual artifacts was aided by simultaneous EOG recordings. Between two and five components were removed for each subject.

### 2.13. MEG source reconstruction

The inner skull, scalp, pial, and grey matter/white matter boundary surfaces generated from Freesurfer and the MRI scan were imported into the Brainstorm software [40] and an automated procedure used to align these data to the MNI coordinate system. The midpoint between the pial surface and grey matter/white matter boundary was extracted and downsampled to 10,000 homogeneously spaced vertices to generate a cortical surface of dipole locations using the *iso2mesh* software [41] implemented in Brainstorm. The inner skull surface was similarly downsampled to 500 vertices. These surfaces were then exported to Matlab, where the scalp surface was used to align the structural data with the MEG digitizers. The aligned MEG gradiometers, inner skull surface, and cortical surface were then used to construct a realistic, subject specific, single shell forward model [42]. Dipole orientations were fixed normal to the cortical surface [43, 44] under the assumption that M/EEG signals are primarily generated by the postsynaptic currents in the dendrites of large vertically oriented pyramidal neurons in layers III, V, and VI of the cortex [45].

Exact low resolution electromagnetic tomography (eLORETA) was used to reconstruct source activity [46, 47]. eLORETA is a linear, regularized, weighted minimum-norm inverse solution with exact, zero error localization [46, 37] which has previously been shown to perform well with (parcellated) resting-state data [37, 48], and is suited to study of whole brain synchronization [49, 50].

The cortical surface was aligned to the MEG-optimized reduction [37] of the HCP-MMP atlas (130 cortical ROIs per hemisphere) in Freesurfer. The time-series of each ROI was calculated as the time course of the first principal component of all voxels within the ROI.

### 2.14. MEG based functional connectome

Functional networks were constructed using phase and amplitude measures within two frequency bands (alpha 8-13 Hz; beta 13-30 Hz). For a given frequency band, data was bandpass filtered. We calculated amplitude envelope correlation (AEC) following multivariate orthogonalization ([51]) between pairs of ROIs to construct functional networks. This metric was chosen to quantify amplitude relationships due to its high reliability [52]. AEC based functional connectivity matrices were thresholded based on a graph metric to maintain a balance between integrated and segregated networks. We compared different connection density values with respect to cost-efficiency which is defined as difference between global efficiency and topological cost expressed as density of network, *K* [53]. We calculated density *K* = 0.25 based on this metric for both alpha and beta MEG connectivity matrices by thresholding to retain strongest 75% connections.

### 2.15. 7T MRI data acquisition for Cohort 2

Whole-brain, high-resolution and high-field structural imaging were acquired from all participants in Cohort 2 on a Siemens 7T Magnetom MRI scanner (Siemens Medical Systems, Germany) using a 32 channel head coil (Nova Medical, USA). The MP2RAGE sequence was used [54], which included two MPRAGE acquisitions with different flip angles and inversion times (echo time 2.68 ms, repetition time 6000 ms, first inversion time 800 ms, second inversion time 2700 ms, first flip angle 7°, second flip angle 5°, voxel size 0.65 × 0.65 × 0.65 mm^3^). To correct for the RF transmit field 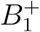, whole-brain 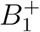 images were acquired using the saturation-prepared with 2 rapid gradient echoes (SA2RAGE) sequence [55] (echo time 1.16 ms, repetition time 2400 ms, first inversion time 540 ms, second inversion time 1800 ms, first flip angle 4°, second flip angle 11°, voxel size 3.25 × 3.25 × 3 *mm*^3^).

### 2.16. 7T MRI data preprocessing and quantitative R1 mapping for Cohort 2

The MP2RAGE sequence generated two images at the first (INV1) and second (INV2) inversion times. For each participant, after an online linear interpolation procedure on INV1 and INV2 maps [54], we obtained a quantitative T1 map that were corrected from proton density contrast, T2* contrast and RF receive field 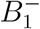 [56]. By combining normalized INV1 and INV2 images, we also obtained a T1-weighted image from the same sequence.

The 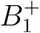 map from the SA2RAGE was registered to individual participant’s INV2 map using the linear registration function FLIRT in FSL [57]. The co-registered 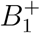 map was then used to correct residual *B*_1_ homogeneities in the T1-weighted and T1 maps [58]. Because the INV2 image has a better contrast between brain tissues and the skull, we used the brain extraction function BET in FSL to obtain a skull-stripped brain mark from the INV2 image. Next, as in previous studies [59, 60], the Java Image Science Toolkit [61, RRID: SCR 008887] and the CBS High-Res Brain Processing tools (RRID: SCR 009452) of the MIPAV platform [62, RRID: SCR 007371] took the T1-weighted image and the brain mark as inputs and generated probabilistic maps of the intracranial dura and arteries. These probabilistic maps were thresholded and used as a mask to remove the dura and arteries in the T1 and T1-weighted images. Finally, for each participant, we calculated a quantitative R1 map (i.e., the longitudinal relaxation rate, *R*1 = 1/*T*1) from the bias-corrected, brain-extracted T1 image, which is sensitive to cortical myelination [63, 64, 65]. A threshold of maximum voxel intensity was applied to the R1 map to discard voxels with large artifacts.

### 2.17. Cortical reconstruction and parcellation of R1 maps for Cohort 2

We projected T1w images on brain surface by using *recon-all* from Freesurfer (6.0.0) in two steps by skipping the skull stripping with *hires* option to keep original high resolution of T1w images. We registered HCP-MMP atlas onto the native T1w space for each subject. After projecting R1 volume on the surface layer at depth 50% between white matter and pial surface with *mr_vol2surf*, mean values on each subparcellated HCP-MMP ROI were calculated by *mri_segstats*.

### 2.18. Intersubject Variation

Intersubject variation (ISV) is calculated for structural and functional connectomes. Consider connectivity matrix *S*_(*r*×*r*)_ where *r* denotes number of ROIs on subparcellated HCP-MMP atlas. We calculated cosine distance (CD) between pairs of subjects as follows:

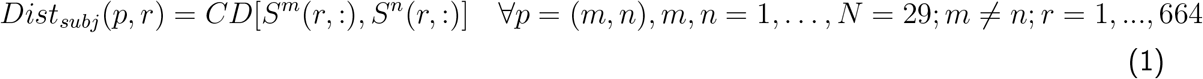

where *S^m^*(*r*, :) denotes the *r*th row of *m*th subject’s connectivity matrix. Cosine distance between two vectors *x* and *y* is defined as:

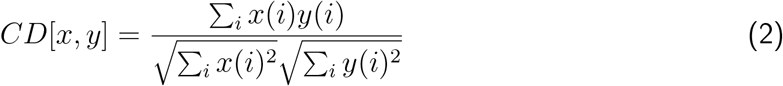

ISV map is calculated from distance maps by averaging across all subject pairs:

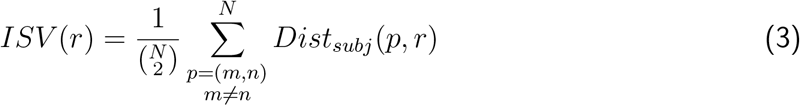

To eliminate the effect of ROI size on intersubject differences, we regressed out mean of ROI sizes across subjects from ISV. We calculated mean ISV on HCP clusters as defined in [11]. Confidence intervals for clusters were calculated based on 5000 bootstrap samples on *Dist_subj_* matrices.

#### 2.18.1. Intersession Variation

For MEG data, we regressed out intersession variance from intersubject variance to remove confounding effects of measurement noise, head motion and low SNR.

Intersession variation for two sessions of MEG data is calculated similar to ISV. In this case cosine distance is calculated between sessions. Consider connectivity matrix *S_r×r_*:

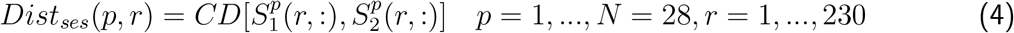

where 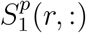 is the *r*th row of *p*th subject’s first session connectivity matrix.

Then, intersession variance is regressed out from each pairwise distance pairs and averaged across pairs:

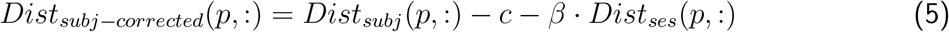

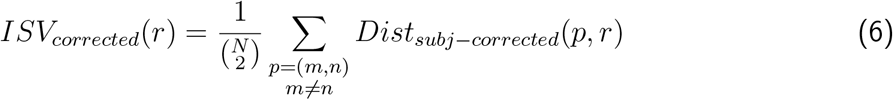

Note that ISV for MEG is calculated for two sessions separately and after correcting for intersession variance, ISV of two sessions are averaged.

### 2.19. Comparison of Imaging Modalities

We compared intersubject variability in structural connectome with functional connectomes to analyse consistency of variability across modalities. For this analysis, after regressing out ROI size from each row of *Dist* matrix in (1), mean of HCP-MMP clusters were found for each subject pairs. Spearman rank correlations were calculated between columns of corrected *Dist* matrix. To assess significancy of correlation between modalities, we used permutation based maximum FWE statistics [66] with 10000 permutation at *p* < 0.05. To avoid effect of intersession variance on MEG *Dist* matrix, we regressed out intersession variance calculated for two subjects from each row of *Dist* matrix. Since MEG data was consisted of 28 subjects, we calculated correlation between 378 subjects pairs for MEG related comparisons and 406 subject pairs for sc-ISV and fc-ISV comparisons.

## 3. Results

### 3.1. Inter subject variability of functional and structural connectivity is not uniform across the brain

We calculated inter subject variability of structural and functional connectomes based on dMRI, fMRI and MEG data from cohort 1. Intersubject variability (ISV) is defined as the average cosine distance between connectivity profile of subject pairs per ROI as in Section 2.18. ROIs were represented on cortical surface parcellations from a recent multimodal HCP atlas [11]. To avoid confounding effect of areal size differences between ROIs, we further subparcellated this atlas into approximately equal sized regions and regressed out areal differences in all computations if it still existed.

Cortical areas in original HCP-MMP atlas were grouped into 22 clusters based on several criteria including neighborship and common functional properties [11]. We quantified average intersubject variability on predefined clusters for structural and functional connectomes.

Structural connectome was constructed based on probability of connections between two regions assessed via tractography (see Section 2.7). Structural intersubject variance (sc-ISV) was calculated on structural connectome (Fig. 2A). We found that sc-ISV was higher in frontoparietal cortices whereas lower in somatomotor, auditory and at the baseline in visual cortices (Fig. 2B-C). Dorsolateral prefrontal cortex (DLPC) and parietal cortex clusters showed the highest sc-ISV in comparison to lowest variability in motor cortex and auditory cortex clusters. Level of variability differed within visual clusters.

**Figure 1:**
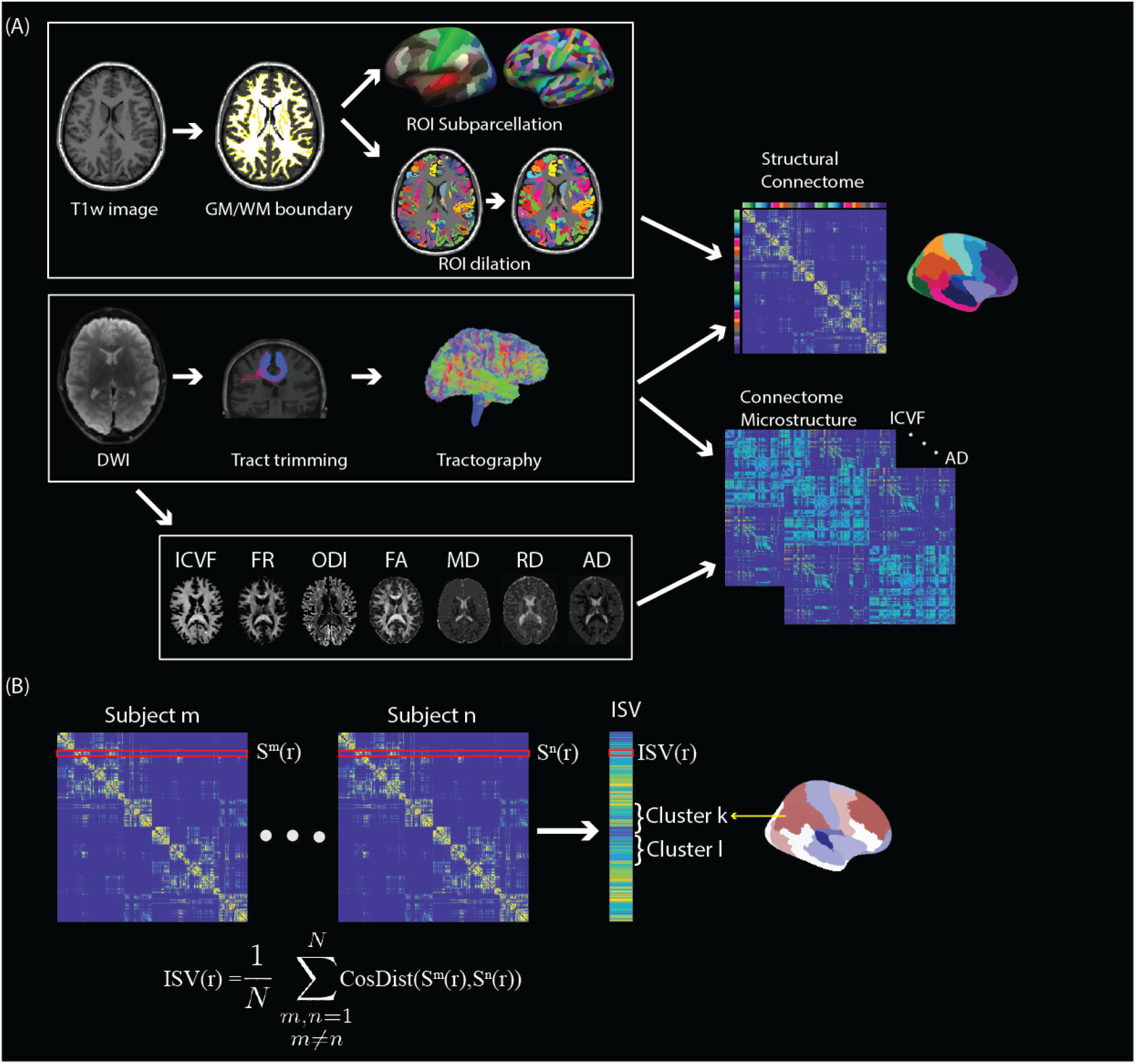
Workflow of structural intersubject variation. **A**, After preprocessing of T1w images, white matter/gray matter boundary (WM/GM) was extracted. HCP-MMP atlas was subparcellated into 664 regions, registered to the individual space and sampled into volume. Cortical parcellation volume was dilated. After preprocessing of DWI, region to region tractography was performed on dilated WM/GM boundary ROIs. Outlier streamlines were trimmed with clustering technique. Finally, structural connectome was generated based on tractography. Microstructural measures from DTI, CHARMED and NODDI models were calculated and sampled along streamlines to form connectomes based on microstructure. **B** Intersubject variance (ISV) per region was defined as the average cosine distance between structural/functional connectivities across subjects. We further grouped ROIs into clusters to compare multimodal ISV.

**Figure 2:**
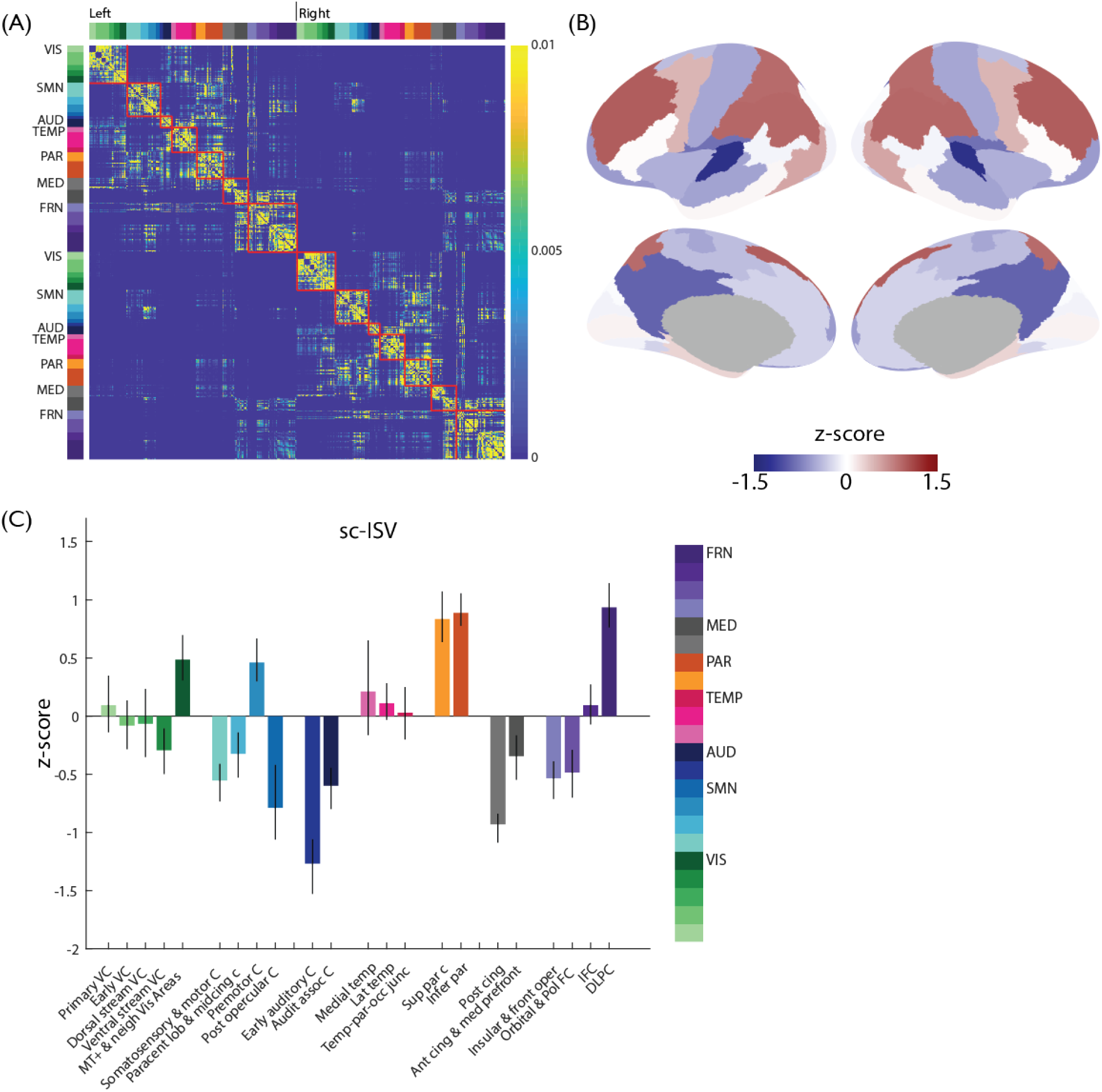
Intersubject variance based on structural connectome (sc-ISV). **A**, Structural connectome of group average. ROIs are grouped as 22 clusters. Clusters in closer spatial location are denoted by similar colors. Borders of groups are marked in connectivity matrix. Colorbar denotes probability of connection calculated by normalizing each row by sum of row itself. **B**, sc-ISV is converted to zscore, averaged across clusters and plotted on cortical surface. sc-ISV higher than mean is denoted by red and lower than mean is denoted by blue. **C**, sc-ISV across clusters. Error bars denote the 95 % confidence intervals. VIS, visual areas; SMN, somatomotor and somatosensory areas; AUD, Auditory areas; TEMP, temporal areas; PAR, parietal areas; MED, medial areas; FRN, frontal areas

To quantify ISV on functional connectivity matrices (fc-ISV) and compare with sc-ISV, we replicated the analysis in [6]. Functional connectivity matrix was obtained by Pearson correlations of fMRI-BOLD signal between subparcellated ROIs and thresholding to maintain the strongest connections (see Section 2.11). We found highest individual variability in frontal and temporal cortices and lowest variability in somatosensory and visual cortices (Fig. 3B). All visual cortex clusters, somatosensory and motor cortices, post opercular, early auditory cortices demonstrated lower fc-ISV than the average. Temporal and frontal clusters had higher fc-ISV (Fig. 3C).

**Figure 3:**
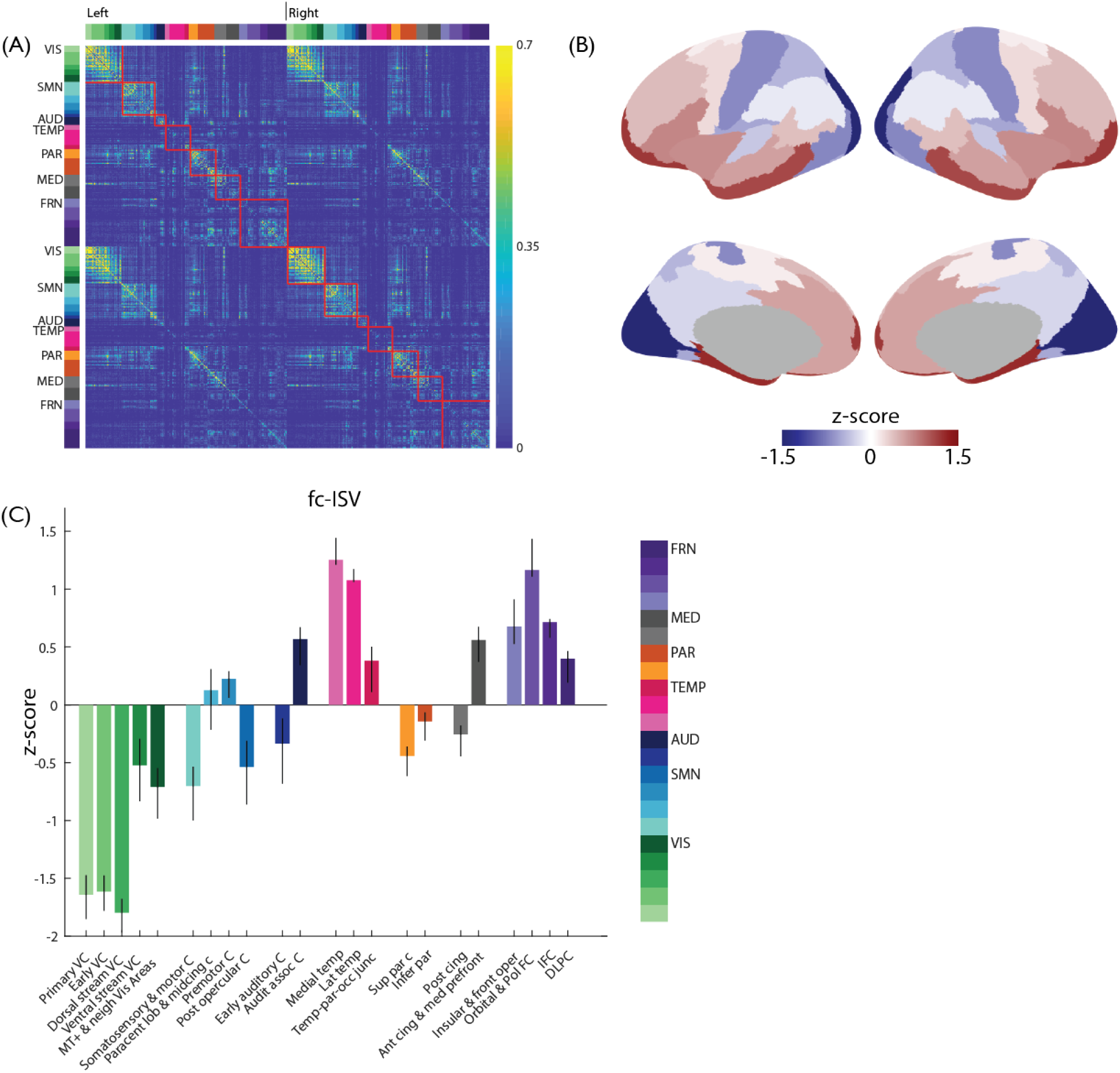
Intersubject variance based on fMRI functional connectome (fc-ISV). **A**, Functional connectome of group average. ROIs are grouped as 22 clusters. Clusters in closer spatial location are denoted by similar colors. Borders of groups are marked in connectivity matrix. Colorbar denotes Pearson correlation between BOLD time series of ROIs. **B**, fc-ISV is converted to zscore, averaged across clusters and plotted on cortical surface. fc-ISV higher than mean is denoted by red and lower than mean is denoted by blue. **C**, fc-ISV across clusters. Error bars denote the 95 % confidence intervals. VIS, visual areas; SMN, somatomotor and somatosensory areas; AUD, Auditory areas; TEMP, temporal areas; PAR, parietal areas; MED, medial areas; FRN, frontal areas

MEG derived functional connectivity quantifies the fast temporal fluctuations of networks that may not be captured by fMRI due to slow hemodynamic response [67, 68]. We calculated MEG connectivity matrix by correlating power envelopes of neural oscillatory signal from downsampled HCP-MMP atlas (see Section 2.14). meg-ISV was calculated for alpha (8-13 Hz) and beta bands (13-30 Hz). Since MEG networks are highly variable within subjects, we regressed out inter-session variance from inter-subject variance [52] (see Section 2.18.1).

meg-ISV was characterized by high variability in frontal-parietal clusters in both bands (Fig. 4C-D). Alpha band meg-ISV demonstrated low variability in all visual clusters in comparison to low variability in all motor clusters in beta band. Temporal clusters had opposite degree of variability in alpha and beta band (Fig. 4E-F).

**Figure 4:**
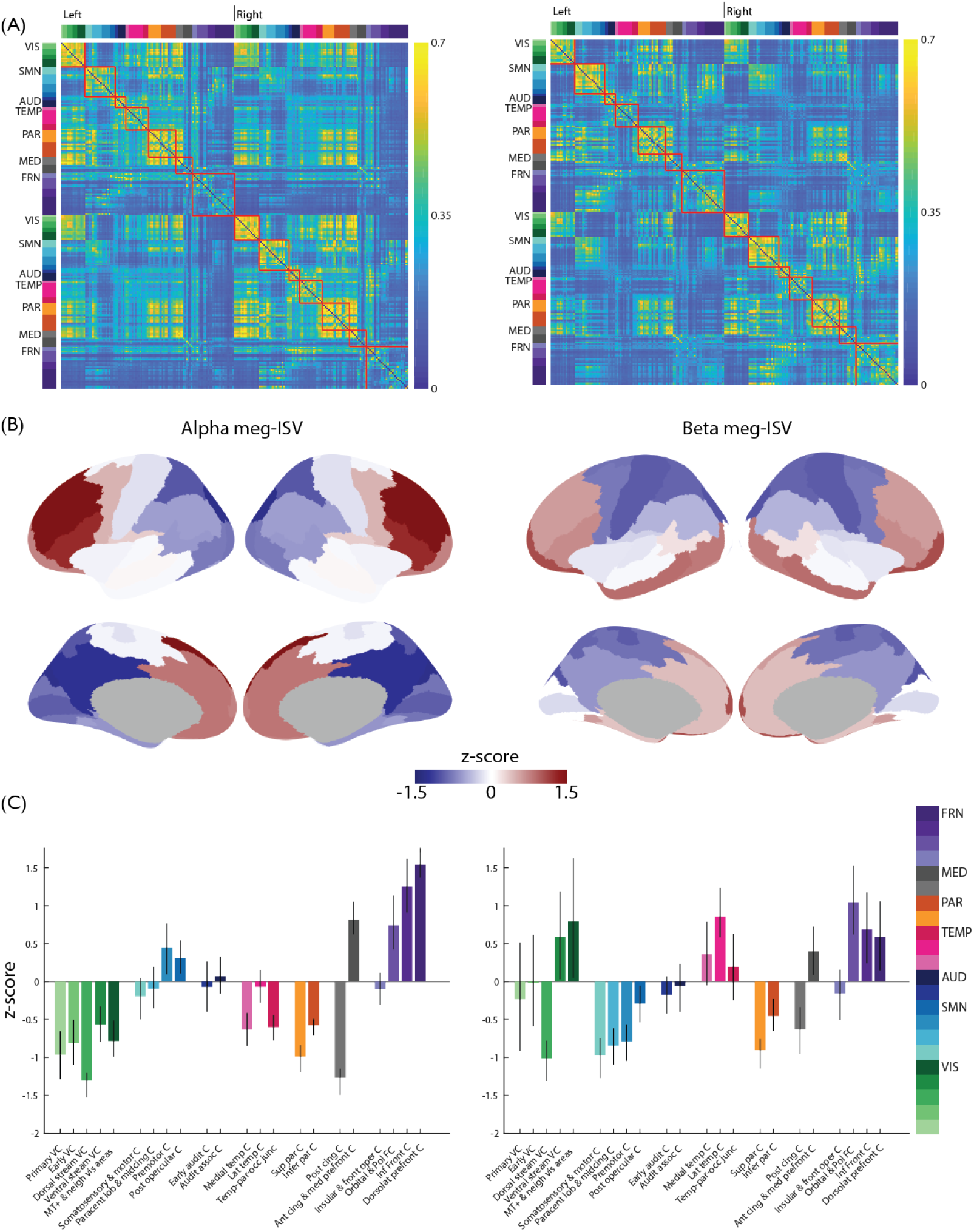
Intersubject variance based on MEG functional connectome (meg-ISV). **A**, Functional connectome of group average. ROIs are grouped as 22 clusters. Clusters in closer spatial location are denoted by similar colors. Borders of groups are marked in connectivity matrix. Colorbar denotes Pearson correlation between amplitude envelope correlation of ROIs. Left alpha band, right is beta band functional connectome **B**, meg-ISV is converted to zscore, averaged across clusters and plotted on cortical surface. meg-ISV higher than mean is denoted by red and lower than mean is denoted by blue. **C**, meg-ISV across clusters. Error bars denote the 95% confidence intervals. VIS, visual areas; SMN, somatomotor and somatosensory areas; AUD, Auditory areas; TEMP, temporal areas; PAR, parietal areas; MED, medial areas; FRN, frontal areas

Evaluation of ISV on functional and structural connectomes demonstrated non-uniformity across cortical surface in all modalities in varying degrees. Next, we assessed alignment between modalities.

### 3.2. Multimodal correspondence of inter subject variability

To test the hypothesis that brain areas have consistent variability across individuals in their structural and functional connectivity, we calculated Spearman rank correlation of distances in SC, FC and MEG connectivity profiles between subject pairs in each HCP-MMP atlas cluster (see Section 2.19). By this way we could avoid the differences in resolution of atlases used for meg-ISV and other ISVs.

Alignment of sc-ISV and fc-ISV was significant in both sensory and association cortices. In particular, frontal, temporal and parietal clusters demonstrated positive correlation. Ventral stream cluster from visual cortex, premotor and paracentral lobular clusters had positive correlation (Fig. 5).

**Figure 5:**
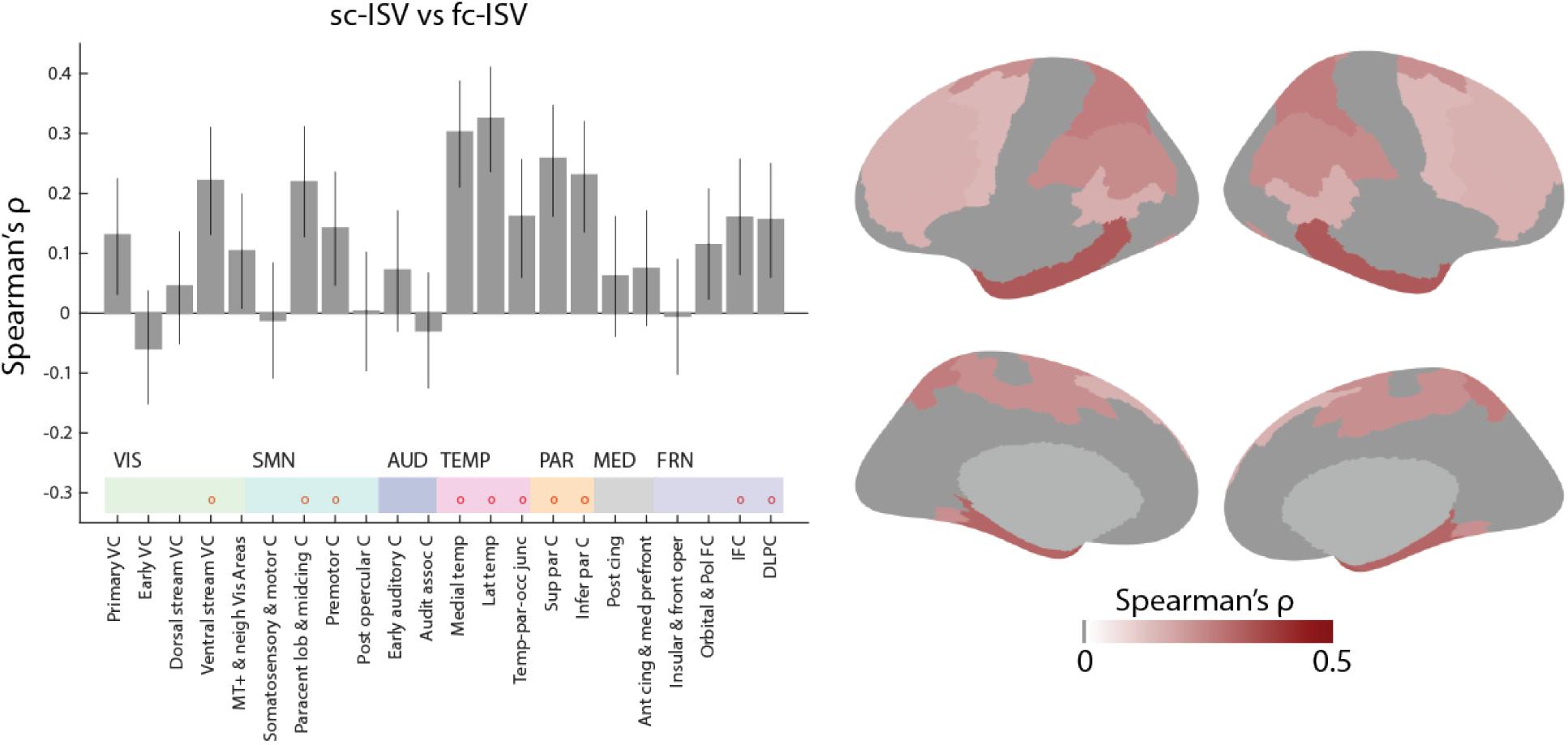
Spearman rank correlations between structural and functional ISV across clusters. The clusters with significant associations with FWE (*p* < 0.05, 10,000 permutations) are denoted with red circles. Error bars denote 95% confidence intervals.

Variability in structural connectome and MEG derived functional connectome was significantly correlated in only a few clusters from frontal, medial, temporal and motor clusters. We observed a wider and higher association between structural variability and meg-ISV in beta band mainly in inferior frontal and parietal clusters. (Fig. 6A). There was a strong alignment in auditory cortex clusters and weak alignment in some visual and somatomotor cortex clusters.

**Figure 6:**
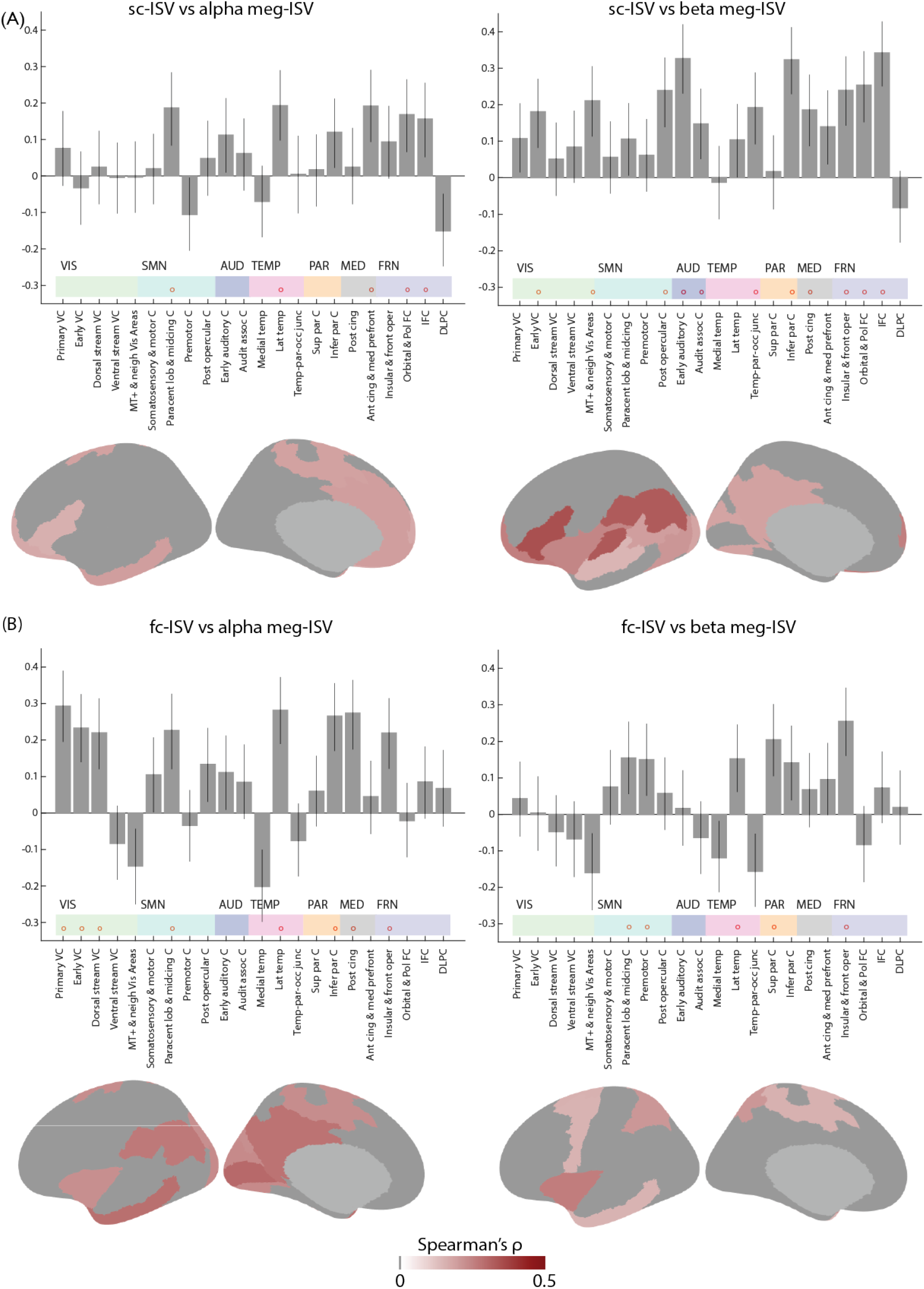
Spearman rank correlations between MEG ISV and structural ISV (**A**) and functional ISV (**B**) across clusters. The clusters with significant associations with FWE (*p* < 0.05, 10,000 permutations) are denoted with red circles. Error bars denote 95% confidence intervals.

Functional variability across subjects assessed with MEG and fMRI had more prominent correlation in alpha than beta band. Variability in functional connectivity of subject pairs were preserved between fMRI and alpha band MEG in visual cortex clusters in contrast to in somatosensory clusters for beta band MEG connectivity (Fig. 6B).

In general, distance between connectivity profiles of subject pairs was preserved across modalities in some HCP clusters.

### 3.3. Highly myelinated cortical regions have lower functional and structural variability

Cortical myelination is a marker for functional specialization and is validated for delineating cortical boundaries [69]. Brain regions with similar cyto- and myeloarchitecture are tended to connect functionally and structurally in animal studies [70, 71, 72]. Longitudinal relaxation rate (R1) is sensitive to myelin enabling quantification of cortical myelin by using non-invasive imaging methods [73, 63].

To test the hypothesis that highly myelinated cortical areas would demonstrate lower variability in their functional and structural connectivity, we correlated R1 maps computed on cohort dataset 2 against sc-ISV, fc-ISV and meg-ISV from cohort dataset 1. It is important to note that to assess correlation between R1 value and meg-ISV, we downsampled R1 value maps from subparcellated HCP-MMP atlas (664 ROIs in total) to MEG optimized HCP-MMP atlas (230 ROIs in total).

R1-derived cortical myelin maps on cohort-2 demonstrated a clear distinction between associative cortices and sensory & motor cortices. Average R1 values across subjects were the highest in primary visual, somatosensory and early auditory clusters whereas lowest in medial and frontal clusters (Fig. 7A-B).

**Figure 7:**
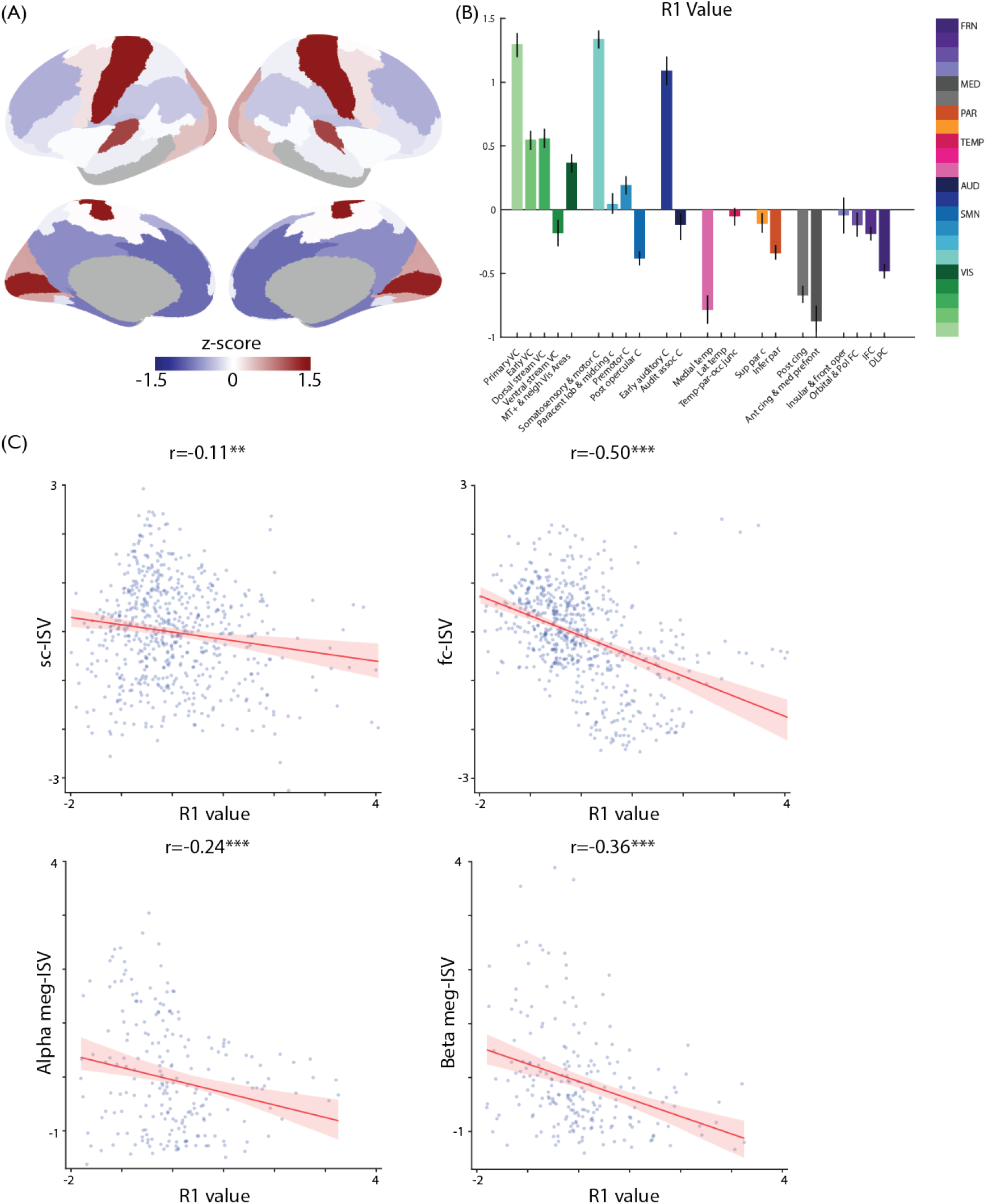
**A**, Group R1 map from Cohort 2 is converted to zscore, averaged across clusters and plotted on cortical surface. R1 value higher than mean is denoted by red and lower than mean is denoted by blue. **B**, R1 value across clusters. Error bars denote the 95% confidence intervals. VIS, visual areas; SMN, somatomotor and somatosensory areas; AUD, Auditory areas;TEMP, temporal areas; PAR, parietal areas; MED, medial areas; FRN, frontal areas. **C**, Group R1 value from Cohort 2 across 664 ROIs correlates with structural, fMRI functional and MEG functional ISV. To correlate with meg-ISV, R1 map is downsampled to MEG optimized HCP-MMP atlas. *r* represents Spearman rank correlation and statistical significance is calculated with a two-sided test permutation test (N=10,000, \*\**P* < 0.0l,\*\*\**P* < 0.001)

**Figure 8:**
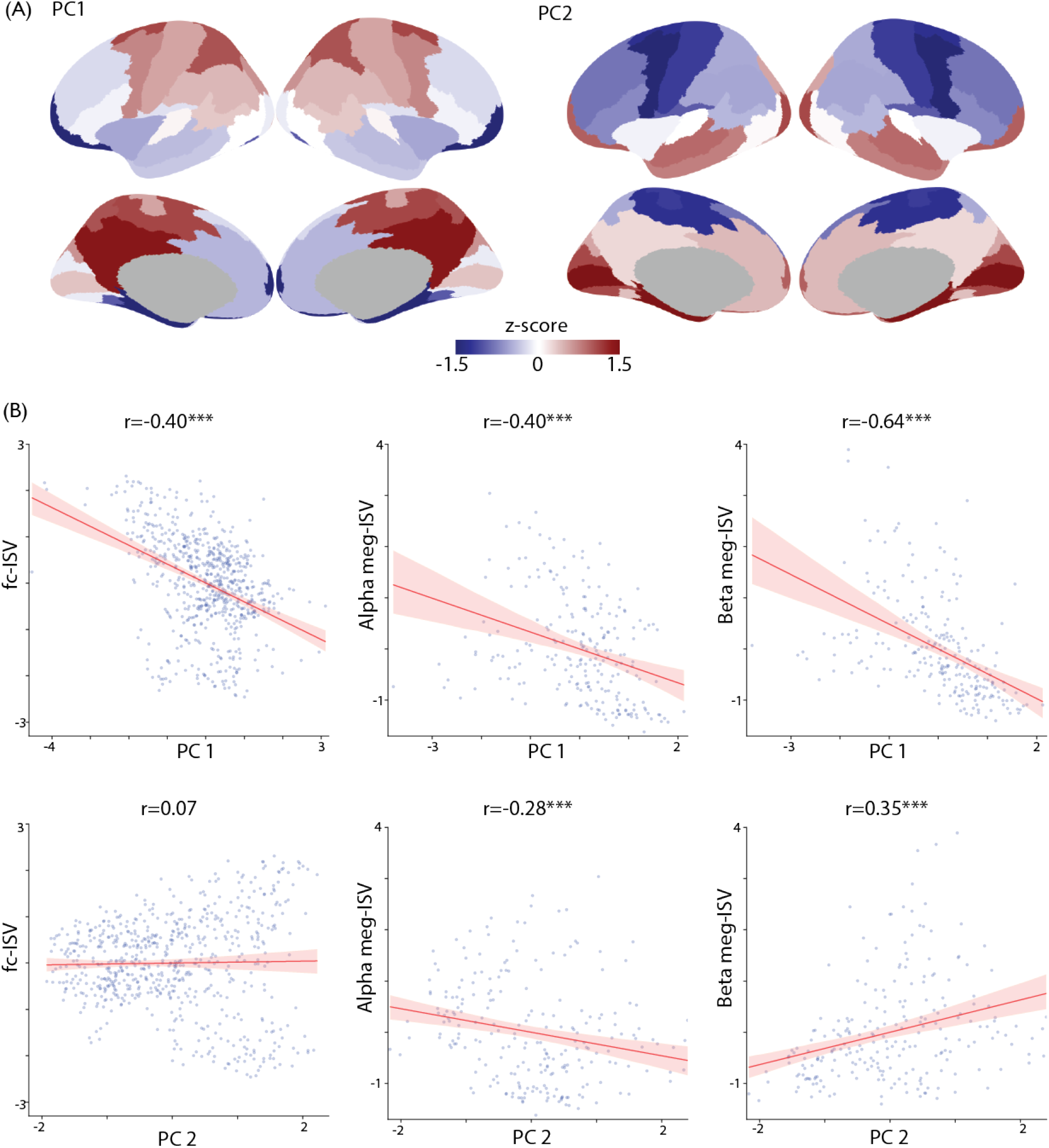
After aggregating connectomes with microstructural measures across subjects, PCA is applied for dimensionality reduction. Two components are retained. **A**, Group PC1 and PC2 maps from Cohort 1 is converted to zscore, averaged across clusters and plotted on cortical surface. Values higher than mean are denoted by red and lower than mean are denoted by blue. **B**, Group PC1 and PC2 from Cohort 2 across 664 ROIs correlate with structural, fMRI functional and MEG functional ISV. To correlate with meg-ISV, PC maps are downsampled to MEG spatial atlas that is composed of 230 ROIs. *r* represents Spearman rank correlation and statistical significance is calculated with a two-sided test permutation test (N=10,000, \*\**P* < 0.01,\*\*\**P* < 0.001)

Structural variability was negatively correlated with R1 myelination map showing a weak anticorrelation (r=-0.11, p=0.009, CI=[−0.18, −0.03]). Functional connectivity variability was negatively associated with myelination map at a moderate level (r=-0.50, p=0.78e-39, CI=[−0.56, −0.43]). Functional connectivity variability assessed by MEG also showed moderate anticorrelation in both bands (alpha: r=-0.24, p=0.0006, CI=[−0.36, −0.10] and beta: r=-0.36, p=1.04e-07, CI=[−0.47, −0.23]) (Fig. 7C).

### 3.4. Functional variability is correlated with white matter microstructure

Microstructural properties of white matter connections assessed with DWI includes degree of anisotropy of water diffusion [74], intracellular and extracellular neurite density [27] and fractional myelin content [28]. Covariation of white matter microstructure and functional connectivity suggests a non-trivial relation [10, 75].

We investigated the effect of white matter microstructure on inter subject functional connectivity variability by applying PCA on the aggregated microstructural connectivity matrices (see Section 2.9).

PCA of microstructural connectome aggregated across subjects reveals two components: (i) one related to anisotropy (PC1) by representing difference between FA, restricted diffusion signal fraction (FR), ICVF and AD from radial diffusivity, ODI and MD (PC1 = (0.5 FA, 0.4 FR, 0.4 ICVF, 0.3 AD) - (0.5 RD, 0.4 ODI, 0.1 MD), explained variance = 53.87 %) (ii) The second component (PC2) related to diffusivity revealed higher contributions from MD and AD with negative contributions from compartmental models (PC2 = (0.6 MD, 0.5 AD, 0.2 FA, 0.2 RD) - (0.4 ICVF, 0.4 ODI, 0.2 FR), explained variance = 36.55 %).

Taking average of anisotropy related component on streamlines that emit from every ROI revealed a non-homogeneous pattern across brain. It was higher in motor, parietal and posterior cingulate cortices. Anisotropy of streamlines connecting frontal and temporal cortices to rest of the brain was moderately low with respect to average. On the other hand, average of diffusivity related component of connections was higher for visual and temporal cortices, medial temporal regions and polar frontal cortices. Regions whose connections showed lowest diffusivity component were motor, parietal and frontal regions.

Inter subject variability of functional connectivity was moderately negatively correlated with the anisotropy related component values of connections (r=-0.4, p=0.22e-26, CI=[−0.47,−0.02]). There was not any significant correlation between functional connectivity variability and PC2 (r=0.07, p=0.07, CI=[−0.33, 0.16]).

We downsampled mean PC1 and PC2 maps to MEG optimized HCP-MMP atlas. Correlation of meg-ISV with PC1 was highly significant for both frequency bands (meg-ISV-alpha: r=-0.40, p=0.9e-9, CI=[−0.51,−0.27], meg-ISV-beta: r=-0.64, p=0, CI=[−0.71,−0.55]). We found a moderate anti-correlation between alpha band MEG-isv and PC2 (r=-0.47, p=0, CI=[−0.38,−0.17]) and positive correlation with beta band MEG-isv and PC2 (r=0.36,p=0.4e-07, CI=[0.24,0.46]).

## 4. Discussion

We demonstrated that regional differences in individual variability of structural and functional connectomes is characterized by higher variability in association cortices and lower variability in sensory and visual cortices. This pattern is consistent across all modalities at varying degrees as shown by significant alignment between functional and structural connectome variabilities at several clusters of brain regions. Regional specificity map of variability is associated with microstructural properties of connections and cortical myelination.

### 4.1. Structural and functional interindividual variability is heterogenous across cortical surface

Regional specificity in functional and structural variability profiles aligns well with evolutionary and developmental expansion pattern across brain [76, 77, 78]. Multimodal association cortices including frontal, temporal, and parietal areas are evolutionarily modern areas and present a higher cortical surface expansion in humans with respect to macaque [79]. Interestingly, high expanding regions are less mature at term gestation functionally and structurally allowing shaping by environmental factors [76, 80, 81] and mature slowly in developmental stages [76]. Regional differences in white matter maturation in later childhood development supports variability in structural connectome [82, 83, 84]. Individual differences in cognitive functions are correlated with high expanding regions during evolution and development [77]. Furthermore, increase in brain size in humans results in greater expansion in distributed frontoparietal cortical networks [85]. Higher functional and structural variability in association cortices observed in all imaging modalities could be explained with regional differences in cortical surface expansion accompanied with local cellular events and late maturation in white matter and synaptic density.

There are several exceptions in structural connectivity variability that do not follow principal axis of variability that extends from unimodal cortices to association cortices. Orbital and polar frontal cortex exhibits lower variability with respect to other frontal areas which might be due to differences in cortical cytoarchitectonic divisions [86] that effect structural connections [71]. Visual regions show more variability in structural connectivity with respect to motor regions. This finding may arise from higher variability in optic radiation with respect to cortico-spinal tract [87, 30].

Variability in functional connectivity assessed by MEG and fMRI is higher than structural connectivity across all regions. Anatomical connections are mainly determined by material and metabolic costs that constrain connection probability between brain regions for optimal topological patterns [88, 89]. This could lead lesser degree of structural variability across individuals without constraining functional connectivity considering divergence between functional and structural connectivities [90, 91, 92].

MEG based functional connectome can reliably extract intrinsic fluctuations between spatially distant brain regions [93, 9, 68, 94, 95]. Though general agreement between interindividual variability across brain in alpha and beta band oscillations, there are differences in variability especially in visual, motor and temporal cortices. Beta band oscillations are dominant feature of sensorimotor system and prominent in interhemispheric connections of motor cortices in resting state [96, 9, 97, 98, 99]. On the other hand, alpha band oscillations reflect neural synchrony across mainly in occipital but also in temporal cortices [100, 94, 101]. Differences in the underlying generators of resting alpha and beta oscillations explain sensitivity of MEG reflecting interindividual variability.

### 4.2. Interindividual variability is consistent across modalities

We did not find significant correlation between structural and functional connectivity variabilities (sc-ISV and fc-ISV) in the global brain scale. When compared connectivity variability across subject pairs, we observed cluster-level alignment is distributed across brain in both association and unimodal cortices with highest in parietal and temporal clusters. This suggests distance between individuals in terms of functional and structural connectivity is preserved in a cluster-specific manner. Presence of alignment in variability might be due to comparable and similar effects of neurobiological and anatomical factors including cortical folding, sulcal depth [6, 76] and late maturation [82, 76] on individual variability.

Variability in structural connectivity and beta band MEG connectivity was significantly correlated mainly in frontal, parietal, and auditory cortex clusters. Superior longitudinal fasciculus (SLF) connecting frontal areas to parietal areas modulates control networks [102, 103]. In particular, individual differences in lateral SLF are associated with synchronization of oscillations during visuo-spatial attention [104, 105] suggesting well alignment between structural and beta band functional variability. Auditory areas are structurally connected with inferior frontal cortex through arcuate fasciculus [106]. This might play role in significant association between structural and MEG beta band connectivity variability. Functional connectivity variability assessed with fMRI and MEG alpha band were significantly related in visual cortices highlighting sensitivity of MEG on reflecting neural synchrony [100, 94].

In comparison, beta band MEG variability and fMRI functional variability correlated mostly in motor cortices as similar in [9].

### 4.3. Intersubject variability is related to cortical myelin content and white matter microstructure

In line with recent research, our findings from R1 quantitative imaging on an independent cohort demonstrated high cortical myelin content in somatosensory, motor, auditory and visual cortices and low myelin content in association cortices including frontal, parietal and temporal areas [69, 107, 64, 63]. Lightly myelinated areas correspond to be responsible for higher cognitive functions and become myelinated later in lifetime [108, 69]. Interestingly, in grey matter, lightly myelinated areas have lower neuronal density, large dendritic arboration, more spine density and more synapses thus more complex intracortical circuitry [109, 69, 110]. On the other hand heavily myelinated cortical regions are thinner with larger number of smaller cells and simpler dendritic trees. Myelin related factors reduce synaptic plasticity by inhibiting neurite growth such as new axonal growth and synapse formation [111, 112]. Furthermore, lightly myelinated frontal and parietal lobes require higher aerobic glycolysis with respect to lightly myelinated anterior insula, temporal pole, anterior cingulate and primary sensory areas [110]. It was speculated that higher aerobic glycolysis and lower cortical myelin content may suit for more adaptable and plastic circuitry characterized in most of association areas. In light of these findings, negative association between R1 map as a proxy for cortical myelin and individual variability in functional and structural connectivity implicates effect of plasticity. Differences in the degree of negative correlation between structural and functional variability may stem from the fact structural connectivity is more sensitive to white matter plasticity with respect to cortical synaptic plasticity [113, 114].

Our findings mark differences in microstructural properties of white matter pathways across brain. Microstructural measures from DTI and biophysical models used in this study are sensitive to local fiber architecture, axonal morphology, and myelin content. Previous studies quantifying tract-specific measurements of tissue microstructure have also found variability both along tract and between tracts [115, 29]. However, our findings are not directly comparable with existing methods since (i) Microstructural metrics were aggregated and projected onto principal components that explain the highest variability (ii) Principal components were averaged across streamlines that connect brain regions. Group averaged first principal component of microstructure that represents anisotropy component demonstrated higher values in primary visual, motor, somatosensory and parietal regions. This might be due to higher homotopic connectivity between those regions [10]. However, it is important to note that our analysis on microstructure is not sensitive in tract level and it is challenging to interpret specific differences between regions. High anisotropy component in this study represents higher neurite and axonal density from compartmental models and lower orientation dispersion that could lead more synchronized network and lower inter subject variability in functional connectome. In line with various studies, we have shown variability in functional connectivity between regions is mediated with microstructural properties of white matter trajectories that connect them [10, 116, 117, 118, 119].

## 5. Conclusion

In conclusion, structural and functional inter subject variability was higher in association cortices and lower in visual, motor and auditory cortices. Variability was preserved across modalities in some clusters of regions. Cortical myelin content and white matter microstructure was related to intersubject variability. Our findings contribute to understanding of individual differences in functional and structural organization of brain and facilitate fingerprinting applications.

## 7. Supplementary

**Figure 9:**
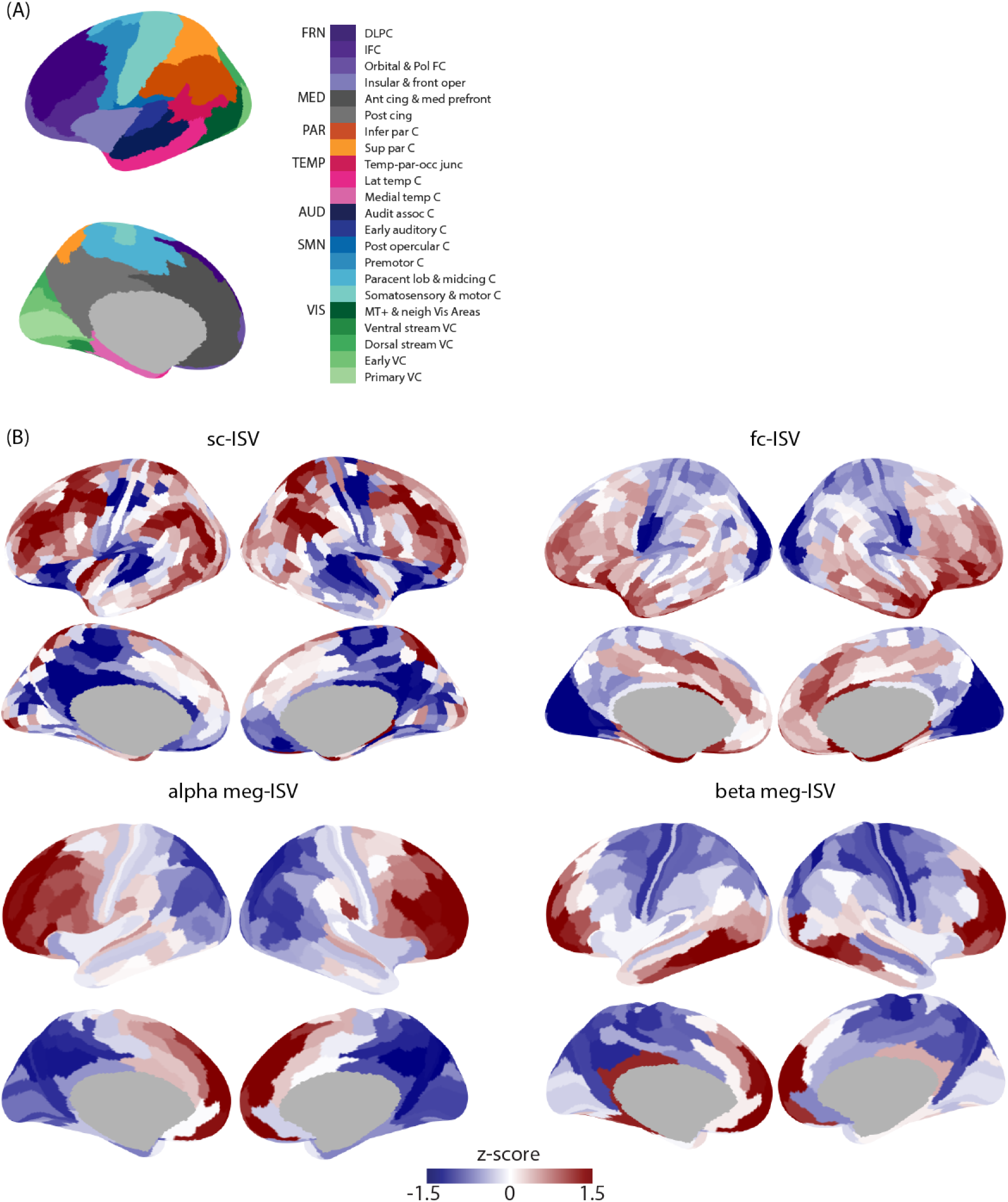
**A**, HCP clusters grouped according to their spatial locations. Clusters in the same group are plotted in similar colors. VIS, visual areas; SMN, somatomotor and somatosensory areas; AUD, Auditory areas;TEMP, temporal areas; PAR, parietal areas; MED, medial areas; FRN, frontal areas. **B** structural and fMRI functional maps are plotted on subparcellated HCP-MMP atlas ROIs. To calculate meg-ISV dowsampled HCP-MMP atlas is used.

**Figure 10:**
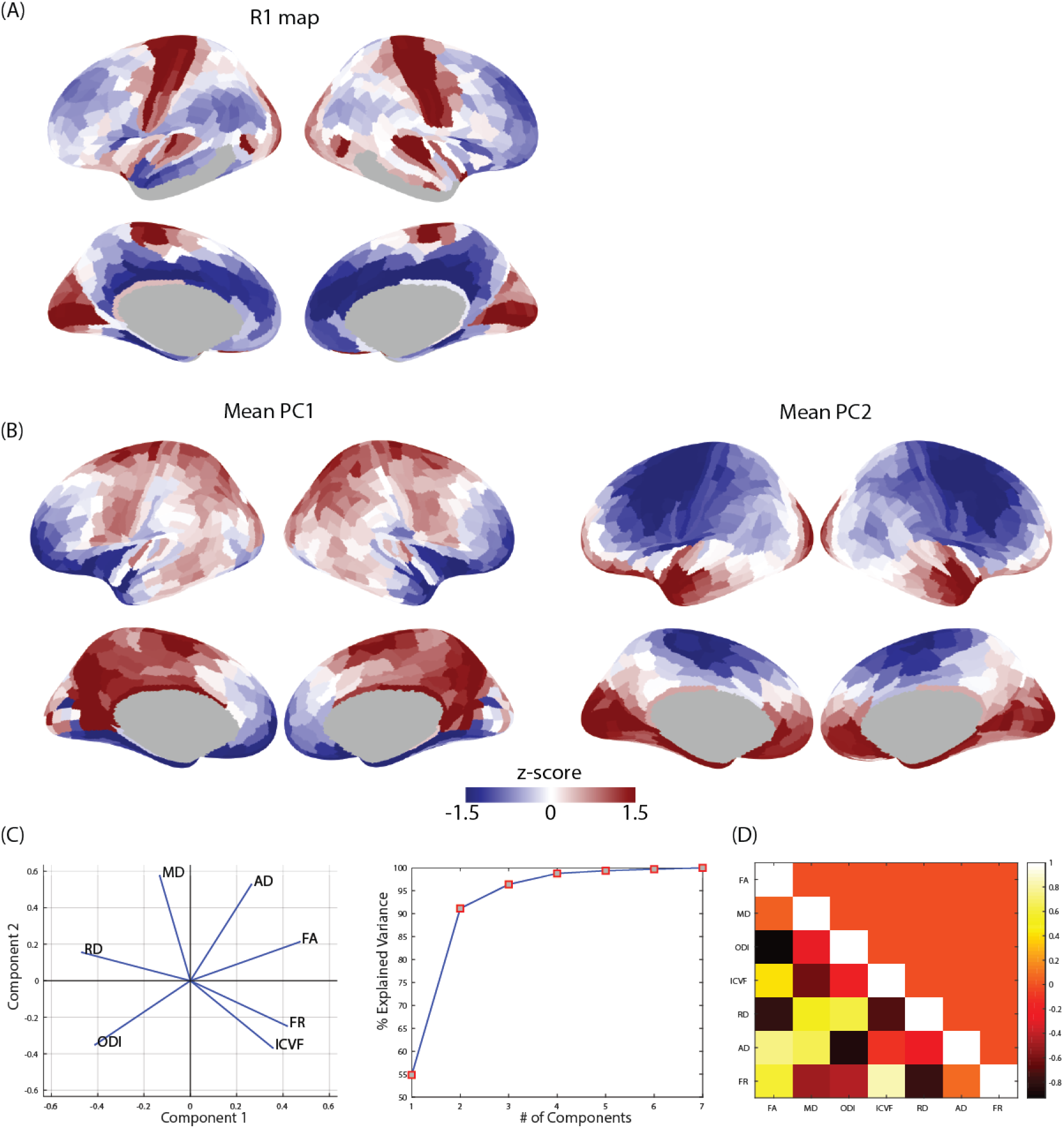
**A**, Group averaged R1 map from Cohort 2. **B** Group averaged first and second principal components of aggregated microstructure connectome from Cohort 2 plotted on HCP-MMP subparcellated atlas. **C** PCA results with 7 microstructural metrics. ICVF, intracellular volume fraction; ODI, orientation dispersion index; FR, restricted water fraction; FA, fractional anisotropy; MD, mean diffusivity; AD, axial diffusivity; RD, radial diffusivity. **D** Correlation between group averaged microstructural metrics across connectome

## Notes

### Competing Interest Statement

The authors have declared no competing interest.

